# Consistent metagenome-derived metrics verify and define bacterial species boundaries

**DOI:** 10.1101/647511

**Authors:** Matthew R. Olm, Alexander Crits-Christoph, Spencer Diamond, Adi Lavy, Paula B. Matheus Carnevali, Jillian F. Banfield

## Abstract

Longstanding questions relate to the existence of naturally distinct bacterial species and genetic approaches to distinguish them. Bacterial genomes in public databases form distinct groups, but these databases are subject to isolation and deposition biases. We compared 5,203 bacterial genomes from 1,457 environmental metagenomic samples to test for distinct clouds of diversity, and evaluated metrics that could be used to define the species boundary. Bacterial genomes from the human gut, soil, and the ocean all exhibited gaps in whole-genome average nucleotide identities (ANI) near the previously suggested species threshold of 95% ANI. While genome-wide ratios of non-synonymous and synonymous nucleotide differences (*dN*/*dS*) decrease until ANI values approach ∼98%, estimates for homologous recombination approached zero at ∼95% ANI, supporting breakdown of recombination due to sequence divergence as a species-forming force. We evaluated 107 genome-based metrics for their ability to distinguish species when full genomes are not recovered. Full length 16S rRNA genes were least useful because they were under-recovered from metagenomes, but many ribosomal proteins displayed both high metagenomic recoverability and species-discrimination power. Taken together, our results verify the existence of sequence-discrete microbial species in metagenome-derived genomes and highlight the usefulness of ribosomal genes for gene-level species discrimination.

## Introduction

A fundamental question of microbiology is whether bacterial genetic diversity exists as a continuum or is divided into distinct clusters (Cohan, 2019, 2002; Shapiro and Polz, 2015). Sequence discrete populations have been identified in public databases (Goris et al., 2007; Konstantinidis and Tiedje, 2005), most recently in a study using ∼90,000 bacterial genomes available in the public NCBI Genome database as of March 2017 (Jain et al., 2018). In these studies, genomes most commonly shared either >97% average nucleotide identity (ANI) or <90%. Based on the observation that relatively few genomes share ANI values near 95%, a bacterial species threshold of 95% ANI was proposed. However it is still unclear whether this pattern is confounded by biases or whether it reflects a true phenomenon across natural environments, as comparison of biased genome sets could form spurious sequence clusters.

Over 75% of the genomes with assigned taxonomy in the NCBI Genome database are from the Proteobacteria and Firmicutes phyla, and over 10% are from the genus *Streptococcus* alone (Jain et al., 2018). Attempts have been made to un-bias this reference genome set when looking for naturally distinct bacterial populations, for example by sampling five genomes from each species with at least five genomes in the database (Jain et al., 2018), but selective cultivation and sequencing cause biases that are difficult to account for. Biases introduced in the databases include sequencing and depositing isolates that meet the expected criteria of target species, and cultivation with selective media that favors certain genotypes and suppresses the growth of alternative ones. Sets of genomes without selection and cultivation biases can be acquired through the direct sequencing of environmental DNA (genome-resolved metagenomics). While metagenomic sequencing does suffer from its own set of biases, including better DNA extraction from gram positive than gram negative bacteria (Albertsen et al., 2015; Guo and Zhang, 2013), it is unlikely that this kind of broad bias would contribute to patterns of species-level sequence groups. The set of genomes that can assembled from metagenomes can also be biased, especially when multiple similar genomes are present in the same sample (Nayfach et al., 2019; Olm et al., 2017). However, these strain-level biases should not affect the ability to resolve species-level groups.

If distinct microbial species do exist, a relatively comprehensive analysis of public data may uncover the roles of recombination and selection in their origin. Several hypotheses have been proposed to explain genetic discontinuities, including a decrease in rates of homologous recombination at the species threshold (Majewski and Cohan, 1999; Vulić et al., 1997), periodic selective events that purge genetic diversity (Gevers et al., 2005), and neutral processes (Wilmes et al., 2009). Computer simulations suggest that both homologous recombination and selection are needed to form genotypic clusters (Fraser et al., 2007), and quantitative population genomic analyses of metagenomics data point to the declining rates of homologous recombination concurrent with sequence divergence as the force behind the clustering (Eppley et al., 2007). While compelling descriptions of speciation have been shown for a limited number of organisms (Cadillo-Quiroz et al., 2012; Shapiro et al., 2012), homologous recombination and selective pressures have not been measured and analyzed at scale across thousands of genomes, nor in direct relation to the proposed 95% ANI species threshold.

A common method of detecting selection is the *dN/dS* ratio, or the ratio of non-synonymous (*dN*) to synonymous (*dS*) nucleotide changes. Deviations from an expected 1:1 ratio can indicate selective pressures, as nonsynonymous mutations usually have a greater impact on phenotype and are thus more likely to be targets of selection than synonymous mutations. Whole genome comparisons of *dN/dS* are nearly always less than 1 (Castillo-Ramírez and Feil, 2013), indicative of purifying selection and likely due to the continuous removal of slightly deleterious nonsynonymous mutations over time.

To understand the species composition of a microbial environment, it is essential to be able to accurately assign sequences to species clusters. While metagenome-assembled genomes (MAGs) can be compared using whole-genome ANI, only a fraction of assembled scaffolds are binned from complex environmental metagenomes. For example, only 24.2% of the reads could be assembled and binned in a rent study of permafrost metagenomes (Woodcroft et al., 2018). More recently, only 36.4% of reads were assembled into binned contigs, and genomes were reconstructed for only ∼23% of the detected bacteria in complex soil metagenomes (Diamond et al., 2019). Absent dramatic improvements in sequencing technologies, complex communities can be more fully characterized through analysis of assembled and unbinned single copy marker genes, for example the 16S rRNA gene, ribosomal genes, or tRNA-ligase genes (Brown et al., 2015; Emerson et al., 2016; Hamilton et al., 2016; Probst et al., 2018). However the accuracy and identity thresholds of these genes for species delineation, as well as their ability to be assembled from metagenomic data, are unknown. It is therefore important to identify marker genes that not only accurately reflect change in species taxonomy and divergence, but also assemble often in metagenomes using common next-generation sequencing technologies.

Here we analyzed thousands of bacterial genomes recovered directly from the sequencing of environmental DNA to test for the existence of a discrete sequence clusters, developed software to estimate the strength of recombination and selection forces between these genomes, and compared over a hundred marker genes for practical species delineation. Discrete sequence clusters were identified in all environments tested, and both estimated recombination rates and genome-wide *dN*/*dS* ratios showed clear patterns in relation to the 95% ANI species threshold. Whole genome ANI methods were compared to various marker gene alignments (including 16S rRNA) for the ability to create species-level groups, and optimal species-delineation thresholds were calculated for each method. Overall our results support the existence of discrete species-level groups for bacteria in the three divergent environments tested, provide sequence-based evidence for the likely evolutionary forces at play, and provide metrics for species delineation in metagenomics studies.

## Materials and Methods

### Preparation of genome sets

Four criteria were used to identify sets of genomes with minimal isolation and selection biases. 1) Genomes must be assembled from DNA extracted directly from the environment without enrichment or culturing. 2) There must be no preference for particular taxa during metagenomic genome binning and/or curation. 3) Genomes must be available from at least 50 samples from the same or similar environments, and there must be at least 1000 genomes total. 4) All genomes must be publically available for download, not just the de-replicated genome set. Many potential metagenomic studies were disqualified based on criteria (3) and (4), leading to the ultimate selection of three genome sets for follow up analysis (Diamond et al., 2019; Olm et al., 2019; Tully et al., 2018). Recent studies involving large-scale genome binning (Parks et al., 2017; Pasolli et al., 2019) could not be included because their pre-de-replication sets included replicate genomes from the same time series’, leading to artificial genome clusters.

In this study, the first analysis set contains 2,178 bacterial genomes from 1,163 premature infant fecal samples, all of which were collected from infants born into the same neonatal intensive care unit (Olm et al., 2019) (**Supplemental Table S1**). These samples are low diversity and Proteobacteria and Firmicutes account for >80% of the bacteria (and for most samples, >90% of the reads could be assigned to genomes). The second set contains 1,166 genomes from the ocean, including Bacteria and Archaea (Tully et al., 2018). The third set contains 1,859 genomes from a meadow soil ecosystem (Diamond et al., 2019) that spans a large number of diverse phylogenetic groups. We also included 4,800 genomes from NCBI RefSeq, accessed February 2018, where we randomly selected 10 genomes from each of the 480 bacterial species with at least 10 genomes.

All publically available genomes available in RefSeq as of February 21, 2018 were downloaded using ncbi-genome-download (https://github.com/kblin/ncbi-genome-download) with the command “ncbi-genome-download --format genbank -p 4 bacteria”. Taxonomy of all genomes was determined using ETE3 (Huerta-Cepas et al., 2016) based on the provided taxonomy ID. A genome set consisting of a subset of the entire RefSeq set was generated to balance taxonomic representation-ten genomes were randomly chosen from of the 480 species in RefSeq that contained at least 10 species, leading to a total of 4,800 genomes. CheckM (Parks et al., 2015) was run on all genome sets and only those with greater than or equal to 70% completeness and less than 5% contamination were retained. All four genome sets are available at https://doi.org/10.6084/m9.figshare.c.4508162.v1.

### Visualization of average nucleotide identity gap

All genomes in each genome set were compared to each other in a pairwise manner using FastANI (Jain et al., 2018), and the genome alignment fraction was calculated by dividing the count of bidirectional fragment mappings by the number of total query fragments. ANI values and genome alignment fraction values were averaged for reciprocal comparisons, and comparisons of genomes to themselves were removed. The density of each combination of ANI and alignment fraction was calculated using scipy.stats.kde (Jones et al., 2001). The density was plotted in a 3-dimensional histogram using matplotlib (Hunter, 2007).

### Calculation of dN/dS and estimated homologous recombination

dRep (Olm et al., 2017) was used to compare each genome set in a pairwise manner on a gene-by-gene basis using the command “dRep dereplicate --S_algorithm goANI -pa 0.8 -con 5 -comp 70”. Briefly, this identifies open reading frames using Prodigal (Hyatt et al., 2010) and compares their nucleic acid sequences using NSimScan (Novichkov et al., 2016). The script “dnds_from_drep.py” was used to calculate the dN/dS ratio among aligned sequences (https://github.com/MrOlm/bacterialEvolutionMetrics). This involves first aligning the amino acid sequences encoded by pairs of genes which at least 70% of the genes aligned with at least 70% sequence identity, and which were reciprocal best hits. Sequences were aligned globally using the BioPython Align. PairwiseAligner (Cock et al., 2009), using a blosum62 substitution matrix, −12 open gap score, and −3 extend gap score. The alignment was then converted into a codon alignment using biopython, and the number of synonymous sites, synonymous substitutions, non-synonymous sites, and nonsynonymous substitutions recorded. Finally, the overall dN/dS was calculated for each genome alignment using the following formula:

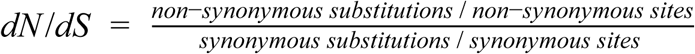

Homologous recombination between genome pairs was calculated as the bias towards identical genes based on the overall ANI between genome pairs, similar to previously described methods (Brito et al., 2016). First, the set of aligned genes was filtered to only those with at least 500 bp aligned. The probability of each gene alignment being identical by chance was determined using the formula:

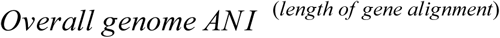

The genome-wide number of expected identical genes was calculated as the sum of the probabilities of each individual gene alignment being the same. The actual number of identical genes for each genome pair was calculated as the number of alignments with a percent identity greater than 99.99%. Finally, the genome-wide bias towards identical genes was calculated using the formula:

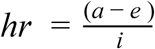

Where *hr* is the estimated degree of homologous recombination, *a* is the actual number of identical genes, *e* is the expected number of identical genes, and *i* is the number of aligned genes with at least 500 bp aligned. Source code is available as python notebooks at https://github.com/MrOlm/bacterialEvolutionMetrics.

### Marker gene identification and clustering

Bacterial single copy genes were identified based on a previously curated set of Hidden-Markov Models (HMMs) for 107 genes expected to be at single copy in all bacterial cells (Albertsen et al., 2013), as accessed on GitHub at the following link on April 10, 2019 https://github.com/MadsAlbertsen/multi-metagenome/blob/master/R.data.generation/essential.hmm. The amino acid sequences of all genomes in all four genome sets were annotated using prodigal in “single” mode (Hyatt et al., 2010) and searched against the single copy gene HMMs using the command “hmmsearch -E .001 --domE .001” (hmmer.org). All hits with scores above the trusted cutoff for each HMM were retained. Nucleic acid sequences for each hit were compared using usearch (Edgar, 2010) with the command “usearch-calc_distmx”. Genes which were identified in less than 85% of genomes in our RefSeq dataset were excluded from further analysis.

16S rRNA genes were identified using SEARCH_16S (Edgar, 2017), with the specific command “usearch -search_16s -bitvec gg97.bitvec”. gg97.bitvec was created using the commands “usearch-makeudb_usearch 97_otus.fasta -wordlength 13” and “usearch -udb2bitvec” based on the Greengenes reference database (as accessed at https://github.com/biocore/qiime-default-reference/blob/master/qiime_default_reference/gg_13_8_otus/rep_set/97_otus.fasta.gz) (DeSantis et al., 2006). Identified 16S rRNA genes were aligned to each other using Mothur (Schloss et al., 2009) with RDP release 11, update 5 (Cole et al., 2014) used as the template. Distance matrices for 16S genes were calculated using the Mothur command dist.seqs.

Recoverability was calculated based on the total number of gene copies that could be recovered from a given metagenomic assembly for each marker gene. This will be impacted by the assembly algorithm, sequencing technology, and nucleotide sequence being assembled. For each set of metagenome-assembled genomes (MAGs), the number of filtered genomes was set as 100% recoverability. The recoverability of each single copy gene was calculated for each set of MAGs using the following equation:

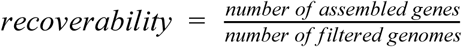

For example, if 100 genomes and 300 sequences of gene *x* were recovered from a set of metagenomes, the recoverability of the gene *x* would be 300%.

### Species delineation scores

Optimal thresholds for species delineation were empirically determined based on pairwise genome distance matrices. For each genome comparison method, all distance thresholds between 80% and 100% were tested, incrementing by 0.1% (80%, 80.1%, 80.2%, etc.). Each pair of genomes at least as similar as the threshold were considered to belong to the same species, and remaining pairs were considered to belong to different species. A species delineation score was calculated for each threshold, and the threshold with the highest score was considered optimal.

Species delineation scores were calculated based the ability to recreate species-level clusters, first as defined by RefSeq taxonomy annotations. A pairwise matrix was established listing each pair of genomes in our RefSeq genome set and whether or not the pair belonged to the same taxonomic species. Recall, precision, and species delineation scores were calculated for each genome set clustering as follows:

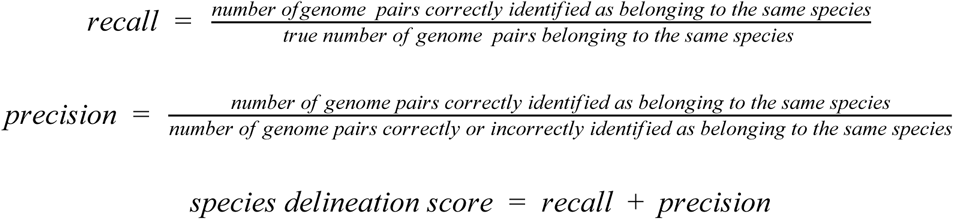

Species delineation scores were also calculated based on the ability to recreate genome clusters as defined by 95% genome-wide ANI similarity (calculated using FastANI). Delineation scores were calculated for all thresholds as described above, and the optimal threshold was defined as the threshold with the highest average delineation score among the three MAG genome sets. Implementation details are available in python notebooks at https://github.com/MrOlm/bacterialEvolutionMetrics.

## Results

### Discrete sequence groups exist in all analyzed genome sets

Sets of microbial genomes without the selection biases introduced by isolation were generated from metagenomic studies of three environments: infant fecal samples (Olm et al., 2019), the ocean (Tully et al., 2018), and a meadow soil ecosystem (Diamond et al., 2019)). A taxonomically balanced set of genomes from Refseq was generated by randomly choosing 10 genomes from each of the 480 species in RefSeq with at least 10 genomes (**Supplemental Table S1**). See methods for details. All genomes within each of the four sets were compared to each other in a pairwise manner using the FastANI algorithm (Jain et al., 2018). Discrete sequence groups based on both ANI and genome alignment percentage were found in all genome sets (**Figure 1**). Notably, species identity gaps were even more prominent in genome sets based on MAGs (metagenome assembled genomes) than RefSeq (which mainly consists of cultured isolate genomes). Comparison of RefSeq genomes marked as belonging to the same vs. different bacterial species showed the identity gap is largely consistent with annotated NCBI species taxonomy, and most genome clusters segregate from each other with a cluster boundary at around 95%. Thus, the analysis is consistent with prior suggestions that this cutoff delineates the species boundary. MAGs from the human microbiome were often very similar to each other (>98% ANI), whereas MAG clusters from the ocean included more divergent strain types. In contrast, most of the comparisons involving genomes from soil involved distinct species.

**Figure 1.**
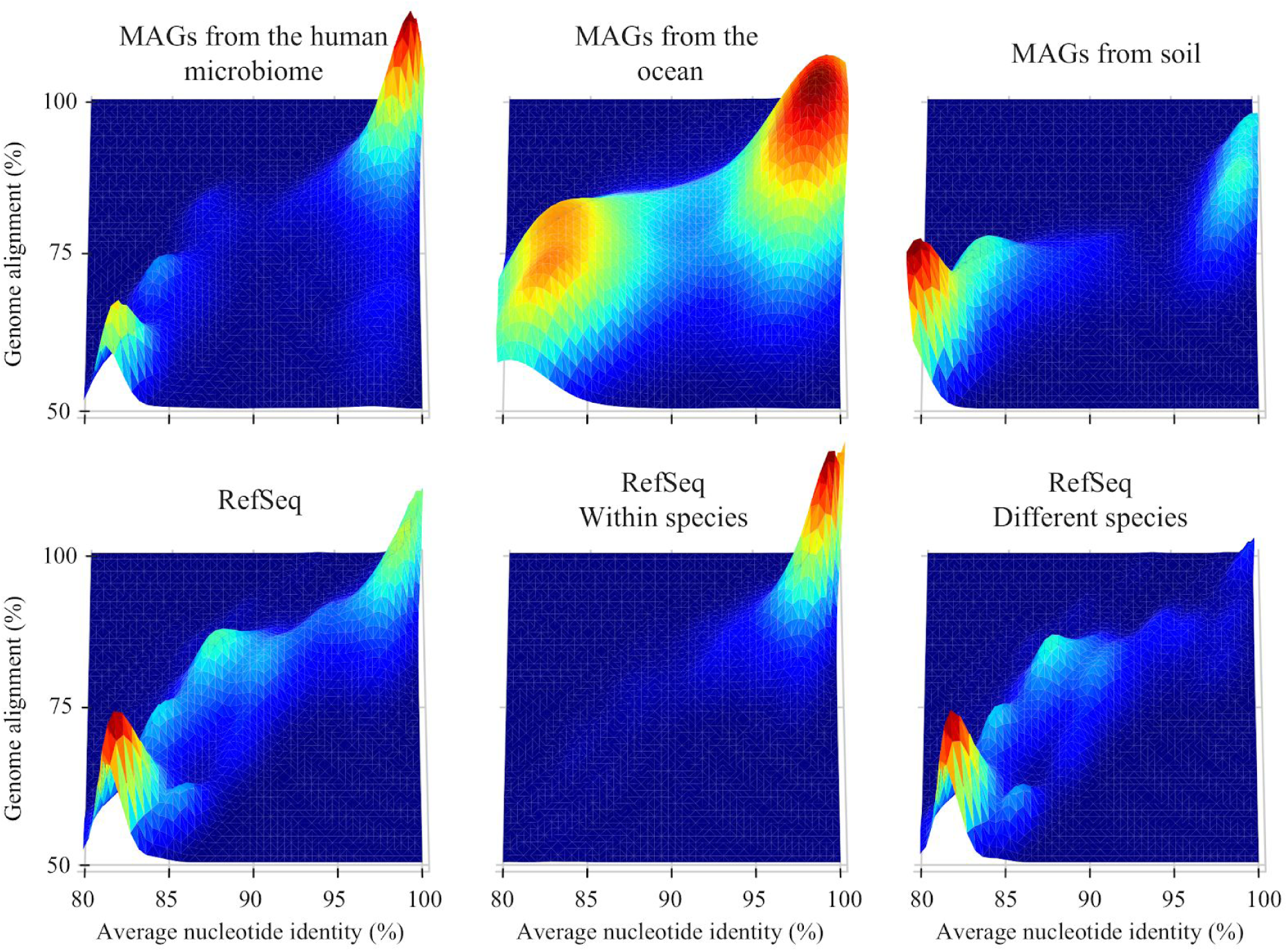
Average nucleotide identity gaps exist near ∼95% ANI in all tested genome sets. Each plot is a histogram of average nucleotide identity and genome alignment percentage values resulting from pairwise comparison within a genome set. Height and color represent density, with higher peaks and hotter colors representing higher numbers of comparisons with that particular ANI and genome alignment percentage. The top row contains three sets of metagenome assembled genomes (MAGs) from different environments. On the bottom row is NCBI RefSeq (rarefied to reduce taxonomic bias; see methods), RefSeq only including comparisons between genomes annotated as the same species, and RefSeq only including comparisons between genomes annotated as different species.

### Gaps in ANI spectra are consistent with measurements of recombination and selection

We next estimated how the evolutionary forces that could lead to discrete sequence clusters change with ANI. An open-source program was written to estimate the rate of homologous recombination and the genome-wide *dN/dS* ratio between each pairwise genome alignment in a rapid and high-throughput manner (see methods for details). Rates of homologous recombination were estimated based on the presence of more identical genes than would be expected by random chance based on the genome-wide ANI, similar to previously described methods (Brito et al., 2016). Genome-wide average *dN/dS* ratios were calculated between pairs of genomes based on a python implementation of the Nei equation (Nei and Gojobori, 1986).

Measurements of estimated homologous recombination and *dN/dS* both followed consistent patterns in relation to the 95% ANI species threshold in all three measured genome sets (**Figure 2**). Estimated homologous recombination rates showed a sharp decline from 100% ANI to around 95% ANI. This relationship could be due to decreasing efficiency of homologous recombination with decreasing sequence similarity.

**Figure 2:**
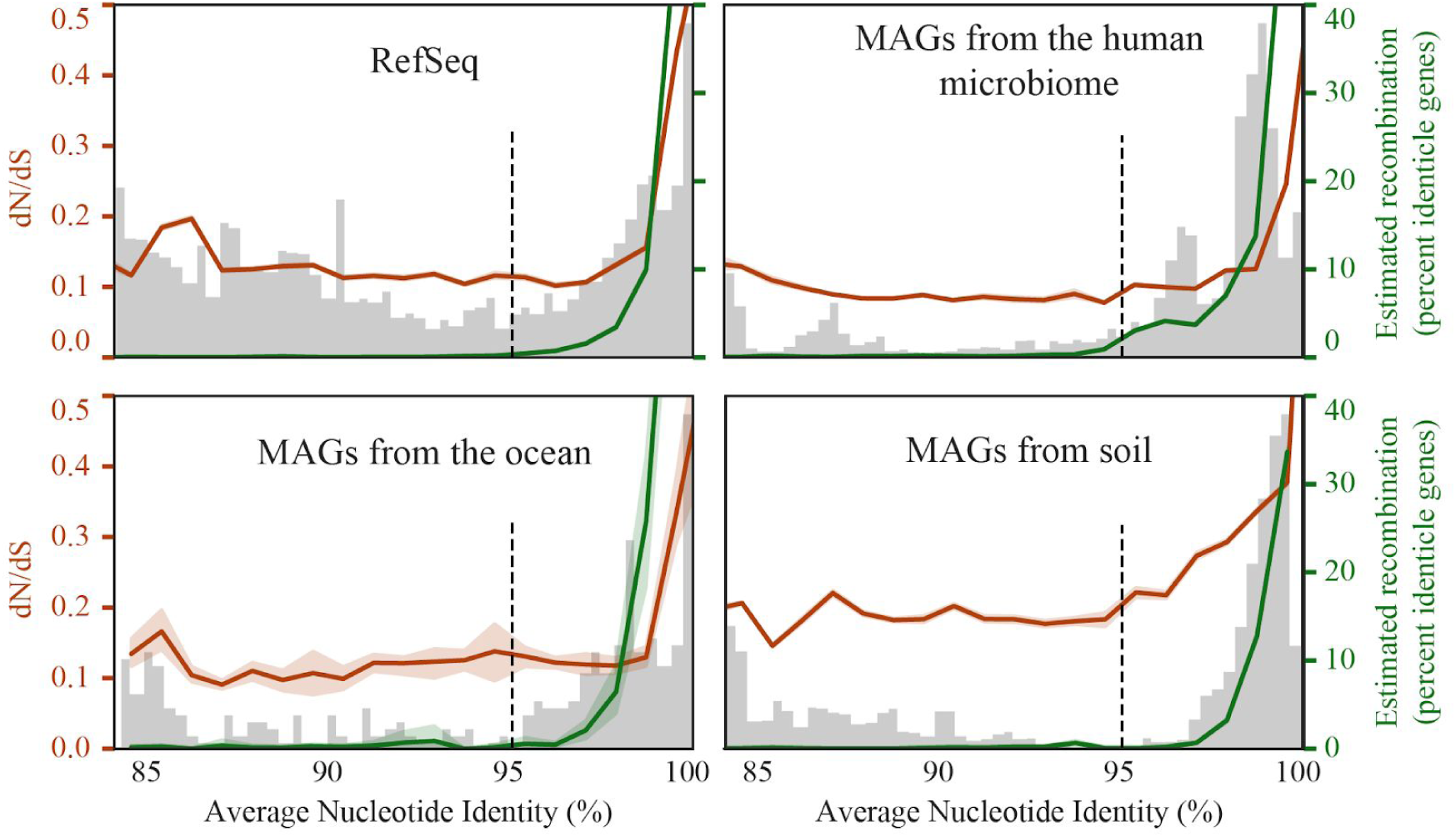
Metrics of recombination and selection follow patterns related to the proposed 95% ANI species threshold. Each plot displays a histogram of ANI values resulting from pairwise comparison within a genome set (light grey bars), the median estimated recombination rate at each ANI level (green line), and the median dN/dS ratio at each ANI level (orange line). A dotted line is drawn at at 95% ANI to mark the commonly proposed threshold for species delineation, and 95% confidence intervals are shown shaded around green and red lines.

All genome-wide *dN/dS* ratios were below 1, as previously observed for whole-genome *dN/dS* comparisons (Rocha et al., 2006). Ratios were highest (∼0.4) between organisms with high sequence similarity and decreased with decreasing ANI, reaching a bottom asymptote of about 0.1 (**Figure 2**). Interestingly, the *dN/dS* asymptote did not tend to occur at 95% ANI, like homologous recombination, but earlier at around 98% ANI. It is well documented that whole-genome *dN/dS* values tend to be higher in recently diverged genomes (i.e., those with high ANI values) (Castillo-Ramírez and Feil, 2013; Rocha et al., 2006), and it is hypothesized that this is because it takes time for purifying selection to purge non-synonymous mutations that are slightly deleterious (nearly neutral). MAG clusters from soil showed a slower decline in *dN/dS* with increasing divergence than was observed in other environments.

### Evaluating marker gene thresholds for bacterial species delineation

To generate an overview of the species composition of an environment, it is necessary to be able to distinguish species from each other. We investigated thresholds for taxonomic species delineation for genome-wide ANI and over 100 marker genes previously identified to occur in single copy in all bacterial genomes (Albertsen et al., 2013) **(Supplemental Table S3; Figure 3)**. These thresholds establish the nucleotide identity shared by genotypes of bacteria considered to be the same species by RefSeq. At values above this threshold genotypes should belong to the same species, and below it, different species. We assigned a score for accuracy of the distinction (precision + recall) using many methods, and found that ANI performs better than any single copy gene (**Figure 3b**). Given that whole genome alignments will generally not be possible for all community members, we also ranked genes for their species discrimination ability. The threshold for the 16S rRNA gene was 99%, identical to that recently reported by Edgar et. al. (Edgar and Valencia, 2018), and significantly higher than the commonly used 97% OTU clustering threshold. The discrimination accuracy for the 16S rRNA gene was among the lowest of the genes considered (**Figure 3b**). Of the genes encoding ribosomal proteins, the gene for ribosomal protein L6 had a high discrimination score and a threshold of <99%, whereas thresholds for tRNA ligase genes and other single copy genes were generally around 97-96% ANI (**Figure 3ab**; **Supplemental Table S2**).

**Figure 3:**
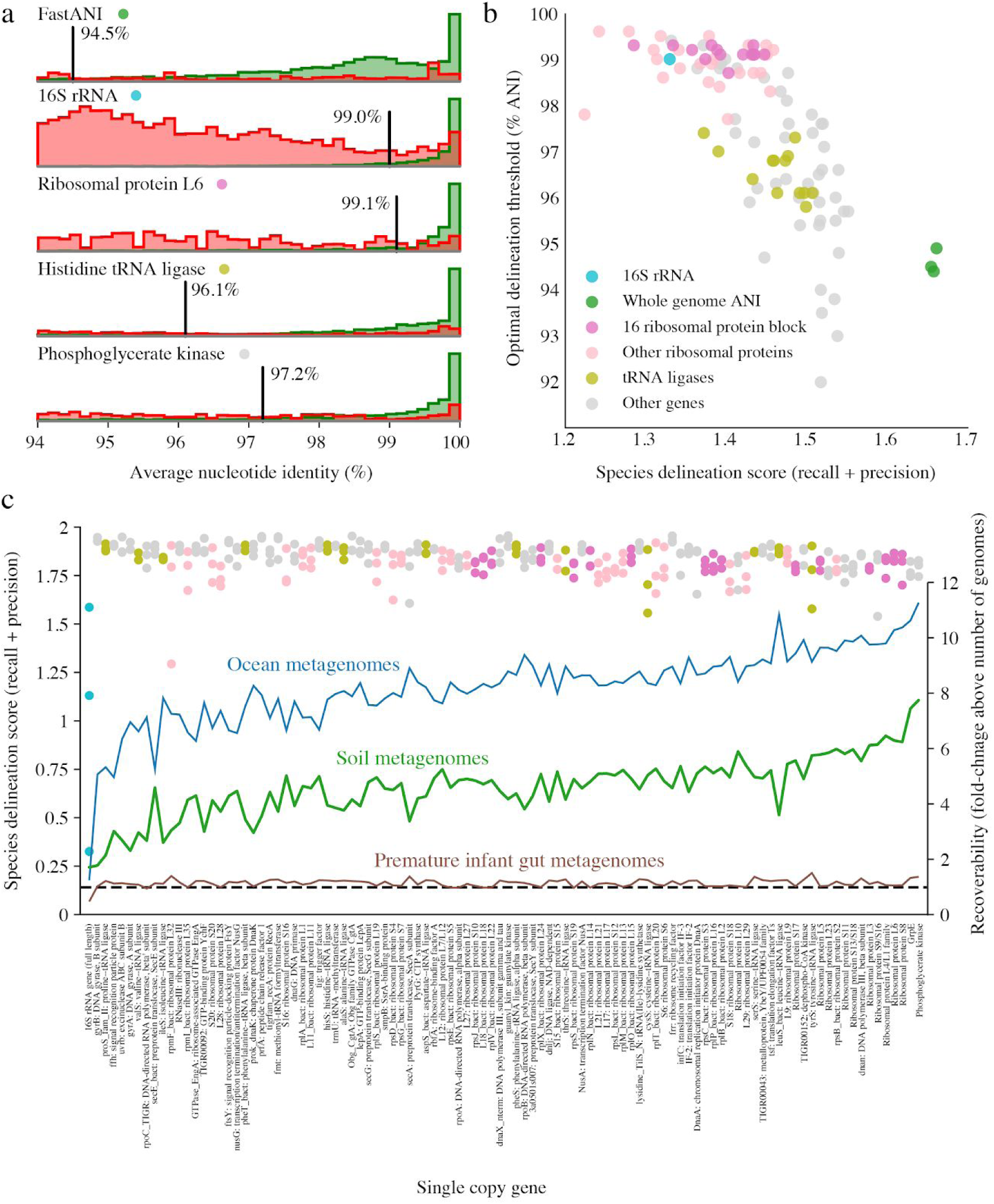
Whole genome alignment outperforms marker genes for species discrimination. (**a**) Histograms of ANI values between bacteria from RefSeq annotated as belonging to the same species (green) or different species (red). Each row is a different method of nucleotide sequence alignment, and vertical black lines indicate the ANI value with the highest species discrimination score for that method. (**b**) Comparison of optimal species discrimination threshold vs. score for reconstruction of species-level clusters from RefSeq. Whole genome comparison algorithms, a 16S rRNA alignment, and single copy gene alignments were tested. (**c**) Accuracy of marker genes to reconstruct species clusters based on 95% ANI whole genome alignments of genomes from metagenomes (dots; left y-axis) and the recoverability of maker genes from metagenomic data from different environments (lines; right y-axis). A horizontal dotted line marks recoverability of 1, meaning an equal number of marker genes and genomes were assembled from the environment.

As RefSeq species classifications are have errors and taxonomic anomalies (Jain et al., 2018), we next compared the ability of marker genes to distinguish genomes that share >95% whole-genome ANI **(Figure 3c)**. The accuracy score for most marker genes was high in all three tested MAG genome sets, with the exception of the 16S rRNA gene which performed poorly **(Figure 3c**). This is likely due to the gene being frequently mis-binned (due to aberrant coverage values resulting from presence in multiple copies) and/or mis-assembled (due to fragmentation caused by its highly conserved regions) in metagenomic data.

It is important that genes used to generate species inventories are easily reconstructed from metagenomes, otherwise species inventories will be incomplete. Thus, we compared the number of marker genes that could be assembled from each dataset to the number of genomes that were assembled and binned from the same dataset, and found that on average five times more ribosomal genes than genomes were recovered (**Figure 3c; Supplemental Table S3**). This was especially true for metagenomes from the ocean and soil, which are complex environments. 16S rRNA genes were recovered much less often than other single copy genes, as has been previously described, but overall there was a wide range in the recoverability of marker genes. Thus, while whole genome comparison methods are most accurate for species-level characterization, many marker genes are good options for species-level marker gene analysis in studies when genomes were not comprehensively recovered. A table listing recommended ANI thresholds based on 95% whole-genome ANI for the 10 single copy genes with the highest recoverability is provided (**Table 1)**, and thresholds for all genes are available in the supplementary material (**Supplemental Table S4**). An open source-program enabling species-level marker gene analysis from metagenomic assemblies is available on GitHub (https://github.com/alexcritschristoph/RPxSuite).

**Table 1.** Species ANI thresholds for the 10 single copy genes with highest recoverability scores

| HMM name | Species ANI Threshold (%) | Species delineation score | Recoverability | Genomes with gene (%) | Genomes with multiple copies (%) | Description |
| --- | --- | --- | --- | --- | --- | --- |
| PGK | 96.5 | 1.8 | 5.12 | 95.2 | 4.8 | Phosphoglycerate kinase |
| GrpE | 96.1 | 1.78 | 4.87 | 96 | 6.8 | GrpE |
| Ribosomal_S8 | 98.3 | 1.8 | 4.44 | 91.9 | 1.6 | Ribosomal protein S8 |
| Ribosomal_L6 | 97.5 | 1.85 | 4.42 | 91.9 | 1.6 | Ribosomal protein L6 |
| Ribosomal_L4 | 98.3 | 1.81 | 4.35 | 91.2 | 1.6 | Ribosomal protein L4/L1 family |
| Ribosomal_S9 | 97.2 | 1.75 | 4.29 | 94.2 | 1.9 | Ribosomal protein S9/S16 |
| Ribosomal_L3 | 98 | 1.79 | 4.26 | 91.2 | 1.5 | Ribosomal protein L3 |
| TIGR00663 | 95.9 | 1.86 | 4.25 | 90.8 | 3.3 | dnan: DNA polymerase III, beta subunit |
| Ribosomal_S13 | 97.9 | 1.79 | 4.24 | 92.8 | 3.2 | Ribosomal protein S13/S18 |
| Ribosomal_S11 | 97.7 | 1.79 | 4.22 | 92.2 | 1.9 | Ribosomal protein S11 |
| 16S | 96.7 | 1.02 | 1.38 | 30.5 | 56.3 | 16S rRNA gene (full length) |

## Discussion

In line with previous studies using reference databases, here we show that bacterial diversity in natural communities is clustered in all three environments studied **(Figure 1)**. Clustering was observed based on both average nucleotide identity and genome alignment fraction (a proxy for shared gene content), estimated rates of horizontal gene transfer fell to near zero at the 95% ANI boundary in all tested environments, and genome-wide *dN/dS* ratios consistently leveled near values of 0.15 at around 98% ANI in most environments **(Figure 2)**. Together these independent metrics support the existence of naturally distinct “bacterial species”.

The observed drop in estimated homologous recombination with decreasing DNA similarity suggests that sequence-dependent homologous recombination is likely a homogenizing force preventing dissolution of bacterial species, in line with previous experimental laboratory studies, computer simulations, (Fraser et al., 2007; Majewski and Cohan, 1999; Vulić et al., 1997), and direct measurements of recombination vs. mutation rates in natural populations (Eppley et al., 2007). Because the drop in *dN/dS* values occurs at ANI significantly above 95%, purifying selection is insufficient to explain the ANI-based species boundary at 95%. This is counter to the stable ecotype model of speciation which posits that species clusters result from series of genetic sweeps (Cohan, 2001; Gevers et al., 2005; Shapiro and Polz, 2015). Taken together, these observations support the notion that bacteria that share >95% ANI recombine often due to shared sequence similarity and that rates approach zero below this value, leading to species divergence and speciation. In combination, the observations support the general applicability of the eukaryotic biological species concept to bacteria (Bobay and Ochman, 2017; Dykhuizen and Green, 1991; Eppley et al., 2007; Rayssiguier et al., 1989).

Given that an increasing number of genomes derive from metagenomic DNA, which do not require culturing or isolation to obtain, a sequence-based method for species delineation is a practical necessity. While thresholds are always prone to exceptions, a genome-wide 95% ANI threshold for species delineation appears to be optimal given the data presented here and previously (Goris et al., 2007; Jain et al., 2018; Konstantinidis and Tiedje, 2005), as well as current species-level taxonomic assignments in NCBI. Here we identified many single copy genes that can act as effective proxies for whole-genome ANI values and are well reconstructed from metagenomes using current technologies **(Figure 3; Table 1)**, and thus are useful for descriptions of microbial communities that are resistant to comprehensive genome recovery. While no tested gene was top ranked in all evaluated metrics, ribosomal proteins S8, L6, and L4 are especially promising candidates given their high recoverability, average species delineation accuracy, and history of use in deep phylogenetic trees (Hug et al., 2016).

## Acknowledgements

This research was supported by the National Institutes of Health (NIH) under award RAI092531A, the Alfred P. Sloan Foundation under grant APSF-2012-10-05, a National Science Foundation Graduate Research Fellowship to M.O. under Grant No. DGE 1106400, m-CAFEs Microbial Community Analysis & Functional Evaluation in Soils, a Project led by Lawrence Berkeley National Laboratory based upon work supported by the U.S. Department of Energy, Office of Science, Office of Biological & Environmental Research under contract number DE-AC02-05CH11231, and Chan Zuckerberg Biohub.

## Data availability

Details of *dN/dS* and homologous recombination analyses are available at https://github.com/MrOlm/bacterialEvolutionMetrics, an open-source program enabling marker-gene analysis from metagenomic data is available at https://github.com/alexcritschristoph/RPxSuite, and nucleotide sequences of genome sets used in this study are available at https://doi.org/10.6084/m9.figshare.c.4508162.v1.

## Author Contributions

M.R.O. and J.B. designed the study; M.R.O. performed the bulk of the metagenomic analyses; A.C.C. contributed to species delineation analysis, S.D., A.L., and P.M.C. contributed to ribosomal proteins analysis; M.R.O. and J.F.B. wrote the manuscript, and all authors contributed to manuscript revisions.

